## Supplementary Information for "The spatial configuration of biotic interactions shapes coexistence-area relationships in an annual plant community"

Note: the code and data to reproduce these results is available at `https://doi.org/10.5281/zenodo.5390313`

### Supplementary Tables

Supplementary Table 1: Species studied and their overall abundance (individuals counted) in the whole study area for each year of observations.

| species | family | 2015 | 2016 | 2017 | 2018 | 2019 |
|---|---|---|---|---|---|---|
| Beta macrocarpa | Amaranthaceae | 136 | 1,230 | 90 | 20 | 1,052 |
| Centaurium tenuiflorum | Gentianaceae | 1,206 | 37 | 1,057 | 522 | 705 |
| Chamaemelum fuscatum | Asteraceae | 5,050 | 5,802 | 2,619 | 0 | 3,989 |
| Chamaemelum mixtum | Asteraceae | 882 | 909 | 208 | 2 | 9 |
| Hordeum marinum | Poaceae | 6,719 | 13,159 | 54,264 | 43,355 | 71,923 |
| Leontodon maroccanus | Asteraceae | 2,485 | 3,448 | 21,657 | 814 | 4,182 |
| Lythrum tribracteatum | Lythraceae | 1,321 | 474 | 676 | 3 | 0 |
| Melilotus elegans | Fabaceae | 313 | 135 | 2 | 0 | 6 |
| Melilotus sulcatus | Fabaceae | 504 | 414 | 36 | 1 | 1,059 |
| Parapholis incurva | Poaceae | 1,239 | 78 | 2,980 | 29 | 7,534 |
| Plantago coronopus | Plantaginaceae | 5,439 | 518 | 2,427 | 1 | 118 |
| Polypogon maritimus | Poaceae | 4,101 | 709 | 5,085 | 315 | 7,332 |
| Polypogon monspeliensis | Poaceae | 442 | 26 | 89 | 1 | 284 |
| Pulicaria paludosa | Asteraceae | 124 | 201 | 2,899 | 746 | 739 |
| Salsola soda | Amaranthaceae | 7,346 | 490 | 1,629 | 4,771 | 791 |
| Scorzonera laciniata | Asteraceae | 881 | 226 | 85 | 1 | 17 |
| Sonchus asper | Asteraceae | 237 | 41 | 0 | 1 | 366 |
| Spergularia rubra | Caryophyllaceae | 146 | 103 | 108 | 0 | 550 |
| Suaeda splendens | Amaranthaceae | 60 | 26 | 0 | 1 | 52 |

Supplementary Table 2: Parameter of power-law fits ($S = cAz$) to the species-area relationships and coexistence-area relationships of Fig. 2. SE = standard error. 2.5%/97.5% = lower and upper bounds of 95% confidence interval for parameter estimates. Statistic and p-values refer to the outcomes of one-sample t-tests. Confidence intervals are obtained with a non-parametric bootstraping procedure. For details on the tests performed, see the function rssoptim in the R package mmSAR.

| year | curve | parameter | estimate | SE | t-value | p-value | 2.5% | 97.5% |
|---|---|---|---|---|---|---|---|---|
| 2015 | heterogeneous | c | 3.47 | 0.91 | 3.81 | 0.01 | 1.65 | 0.91 |
| 2016 | heterogeneous | c | 3.98 | 0.38 | 10.37 | 0.00 | 3.21 | 0.38 |
| 2017 | heterogeneous | c | 1.73 | 0.39 | 4.43 | 0.00 | 0.95 | 0.39 |
| 2018 | heterogeneous | c | 0.64 | 0.20 | 3.15 | 0.02 | 0.23 | 0.20 |
| 2019 | heterogeneous | c | 2.37 | 0.33 | 7.28 | 0.00 | 1.72 | 0.33 |
| 2015 | homogeneous | c | 0.77 | 0.12 | 6.72 | 0.00 | 0.54 | 0.12 |
| 2016 | homogeneous | c | 1.84 | 0.37 | 4.96 | 0.00 | 1.10 | 0.37 |
| 2017 | homogeneous | c | 0.97 | 0.17 | 5.88 | 0.00 | 0.64 | 0.17 |
| 2018 | homogeneous | c | 1.06 | 0.12 | 8.56 | 0.00 | 0.81 | 0.12 |
| 2019 | homogeneous | c | 1.00 | 0.21 | 4.84 | 0.00 | 0.59 | 0.21 |
| 2015 | richness | c | 7.69 | 1.30 | 5.92 | 0.00 | 5.09 | 1.30 |
| 2016 | richness | c | 6.71 | 0.65 | 10.33 | 0.00 | 5.41 | 0.65 |
| 2017 | richness | c | 4.77 | 0.83 | 5.78 | 0.00 | 3.12 | 0.83 |
| 2018 | richness | c | 0.86 | 0.07 | 12.01 | 0.00 | 0.72 | 0.07 |
| 2019 | richness | c | 5.77 | 0.76 | 7.58 | 0.00 | 4.25 | 0.76 |
| 2015 | heterogeneous | z | 0.25 | 0.04 | 5.57 | 0.00 | 0.16 | 0.34 |
| 2016 | heterogeneous | z | 0.22 | 0.02 | 13.41 | 0.00 | 0.19 | 0.25 |
| 2017 | heterogeneous | z | 0.35 | 0.04 | 9.33 | 0.00 | 0.28 | 0.43 |
| 2018 | heterogeneous | z | 0.40 | 0.05 | 7.61 | 0.00 | 0.30 | 0.51 |
| 2019 | heterogeneous | z | 0.31 | 0.02 | 13.43 | 0.00 | 0.26 | 0.36 |
| 2015 | homogeneous | z | 0.46 | 0.02 | 18.62 | 0.00 | 0.41 | 0.51 |
| 2016 | homogeneous | z | 0.34 | 0.03 | 10.16 | 0.00 | 0.28 | 0.41 |
| 2017 | homogeneous | z | 0.44 | 0.03 | 15.59 | 0.00 | 0.38 | 0.50 |
| 2018 | homogeneous | z | 0.29 | 0.02 | 14.73 | 0.00 | 0.25 | 0.33 |
| 2019 | homogeneous | z | 0.45 | 0.03 | 13.01 | 0.00 | 0.38 | 0.51 |
| 2015 | richness | z | 0.15 | 0.03 | 5.05 | 0.00 | 0.09 | 0.20 |
| 2016 | richness | z | 0.17 | 0.02 | 10.04 | 0.00 | 0.13 | 0.20 |
| 2017 | richness | z | 0.20 | 0.03 | 6.93 | 0.00 | 0.14 | 0.26 |
| 2018 | richness | z | 0.45 | 0.01 | 32.66 | 0.00 | 0.42 | 0.48 |
| 2019 | richness | z | 0.18 | 0.02 | 8.09 | 0.00 | 0.14 | 0.23 |

Supplementary Table 3: Coefficients of a logistic regression (with logit link function and N = 171) for the relationship between species abundance (log-transformed) and the classification of species as coexisting or transient. SE = standard error.

| term | estimate | SE | z-value | p-value |
|---|---|---|---|---|
| intercept | 0.62 | 0.45 | 1.36 | 0.18 |
| log(abundance) | -0.18 | 0.07 | -2.63 | 0.009 |

### Supplementary Figures

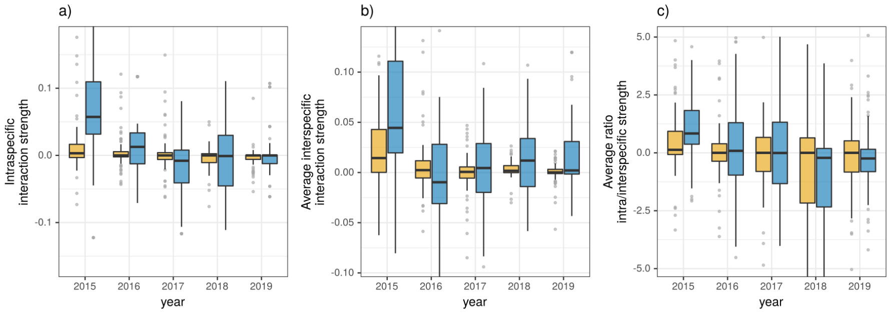


Supplementary Figure 1: Interaction strength distributions across years, for the homogeneous (blue, right-side boxplots) and heterogeneous (yellow, left-side boxplots) parameterizations: (panel a) intraspecific, panel b) average interspecific, panel c) intra/interspecific ratio). In the boxplots, solid black line represents the median of the distribution, the hinges represent the first and third quartiles, and the whiskers extend from the respective hinges to at most 1.5*IQR (interquartile range). Number of observations per year: Panel a) 65,83,66,35,76 (homogeneous), 78,95,77,42,99 (heterogeneous). Panel b) 77,95,76,38,99 (homogeneous), 78,95,76,41,99 (heterogeneous). Panel c) 65,83,65,31,76 (homogeneous), 77,95,76,41,99 (heterogeneous).

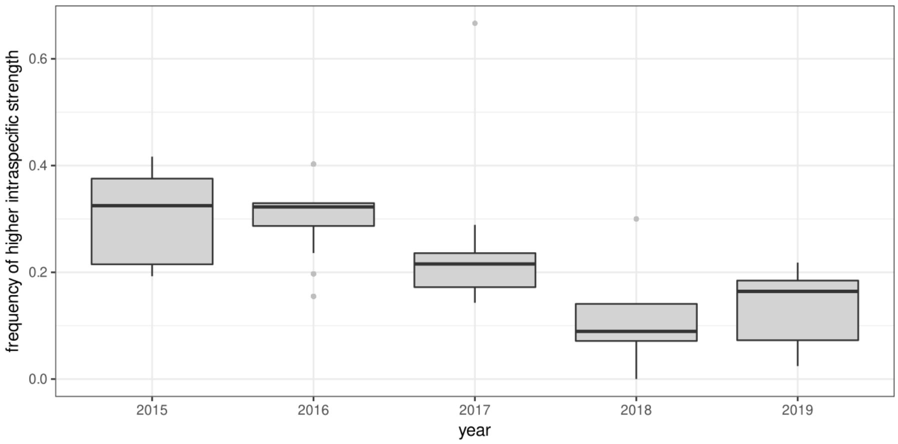


Supplementary Figure 2: Frequency of observing higher intraspecific interaction strengths than interspecific ones, for all observed pairwise combinations of species in each year (N = 820,929,809,248,1053 for 2015,2016,2017,2018,2019 respectively). Results shown for the heterogeneous parameterization. Boxplot summary statistics as in Supplementary Figure 1.

### Supplementary Note 1

In the statistical approach used, we obtain interaction coefficients and intrinsic fecundity rates from a negative binomial model, with plot and year as potential random factors (see main Methods). This model is implemented in a bayesian framework, and in the main analyses we use the means of the posterior distributions for each parameter. To ensure that this reflected the overall trend of the full posterior distributions, and that our results were robust to deviations from these means, we obtained additional coexistence-area curves for each parameterization (homogeneous and heterogeneous, see main Methods) by taking random draws from the posterior distribution of each parameter instead of its mean value. We calculated 100 such replicated curves for each parameterization. Here we draw the obtained coexistence-area curves for each replicate alongside the main ones. The variability associated with these replicates does not deviate qualitatively from the trends obtained with the average posterior values. In particular, we note a higher variability in the curves from the homogeneous parameterization (lower panels of Supplementary Fig. 3) than in the curves from the heterogeneous parameterization (higher panels). This suggests that community-level responses to variations in interaction coefficients or vital rates can be dampened by spatial heterogeneity in such coefficients, e.g. as a result of environmental heterogeneity.

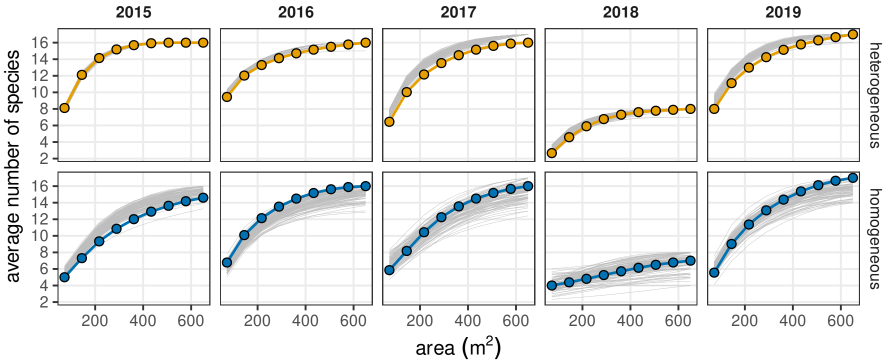


Supplementary Figure 3: Coexistence-area curves in our study system 1) in color, pointed, using the mean of the posterior distributions of the model parameters (interaction coefficients and intrinsic fecundity rates). These are the same curves and follow the same color scheme as in Fig. 2 of the main text. 2) in light grey, obtained from random draws of the posterior distributions of each parameter. For each year and parameterization, 100 curves are represented.

### Supplementary Note 2: Seed survival and germination rates

In order to apply the structural stability framework to the parameters estimated with the Ricker model used in the main analyses, we need to translate the negbin parameters to a form equivalent to a linear Lotka-Volterra model (Saavedra et al. 2017, referenced in main text).

The negative binomial model as applied to annual plants is:

$$\frac{N_{i,t+1}}{N_{i,t}} = (1 - g_i)s_i + g_i F_i \tag{1}$$

where $s_i$ is the survival rate of seeds in the seed bank, $g_i$ is seed germination rate, and $F_i$ is annual fecundity:

$$F_i = e^{\lambda_i - \sum_{j=1}^{n} \alpha_{i,j} N_j} \tag{2}$$

Fortunately, this formulation can be used without further modifications in the structural stability framework. The formula for $F_i$, however, is plugged in the first equation in order to obtain population dynamics, so we need to obtain the feasible equilibrium $N^* > 0$ of the overall model in order to see the equivalence between the annual plant Ricker model parameters and the LV linear model. For the linear LV, this equilibrium reads:

$$r_i = \sum_{j=1}^{n} \alpha_{i,j} N_j \tag{3}$$

We need to derive an equivalent formulation for the negbin annual plant model. As in Appendix S4 of Saavedra et al. (2017), we set

$$(1 - g_i)s_i + g_i e^{\lambda_i - \sum_{j=1}^{n} \alpha_{i,j} N_j} = 1 \tag{4}$$

Reorganizing this equation gives:

$$e^{\lambda_i - \sum_{j=1}^{n} \alpha_{i,j} N_j} = \frac{1 - (1 - g_i)s_i}{g_i}$$

$$\lambda_i - \sum_{j=1}^{n} \alpha_{i,j} N_j = log(\frac{1 - (1 - g_i)s_i}{g_i})$$

$$\sum_{j=1}^{n} \alpha_{i,j} N_j = log(\frac{1 - (1 - g_i)s_i}{g_i}) + \lambda_i$$

so that the equivalence with the linear LV model is:

$$r_i = log(\frac{1 - (1 - g_i)s_i}{g_i}) + \lambda_i \tag{5}$$

This is implemented in the function `growth_rate_equivalence` in the associated code. The interaction coefficients $\alpha_{i,j}$ are not modified. This equivalence shows that seed survival ($s_i$) and seed germination rates ($g_i$) directly influence the estimated growth rates. We performed an additional analysis to explore numerically this influence. In particular, we evaluated to what extent the shape of coexistence-area relationships varied with varying rates. For doing so, we performed additional simulations in which we decreased, increased, or randomly varied each rate by a constant factor: one standard deviation from the distribution of the original samples. This shows whether any directional change exists in response to these increases or decreases, and whether the outcomes are sensitive to random variations in these. Here we show the outcomes of these treatments for the curves of the heterogeneous parameterization. As expected, these variations do not alter the shape of the obtained CARs, with most of them even showing identical curves. Actually, the underlying number of feasible combinations need not be identical in order to display identical curves: slight variations are shown in Supplementary Table 4 in regards to the number of feasible combinations obtained with each treatment. Different spatial combinations of species may clearly give rise to identical CARs. Overall, the variability introduced in the sensitivity analysis of Supplementary Note 2, in which we considered the whole variability in $\lambda$ and $\alpha$ parameters as estimated by the statistical model, is much higher than the variability induced by *in silico* variations of $g$ and $s$. Therefore, in this formulation, variations in seed survival or germination rate have a comparatively negligible influence over the LV-transformed growth rates, and accordingly, on the feasibility patterns of the studied communities. This is, of course, a result contingent on the necessary adaptation of the annual plant model to a linear Lotka-Volterra formulation. Other mathematical strategies to explore coexistence, and the role of these vital rates, may complement this view, but that is outside the scope of this study.

Supplementary Table 4: Number of feasible species combinations each year (summed across plots), in the main analysis and with different modifications of seed survival and germination rates. "Main" refers to the original analysis.

| analysis | 2015 | 2016 | 2017 | 2018 | 2019 |
|---|---|---|---|---|---|
| higher germination rate | 1114 | 688 | 365 | 50 | 474 |
| higher seed survival | 1121 | 687 | 361 | 50 | 471 |
| lower germination rate | 1122 | 675 | 361 | 51 | 469 |
| lower seed survival | 1117 | 675 | 361 | 50 | 472 |
| main | 1124 | 679 | 361 | 50 | 468 |
| random variation in germination rate | 1303 | 743 | 381 | 53 | 499 |
| random variation in seed survival | 1124 | 681 | 361 | 50 | 469 |

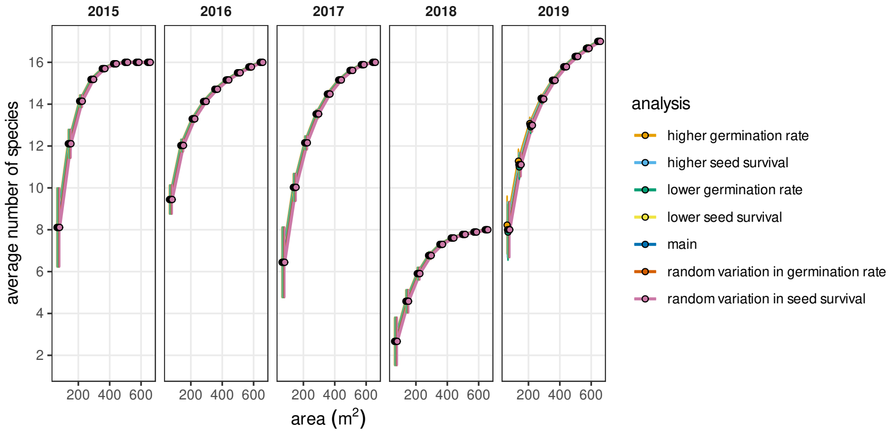


Supplementary Figure 4: Coexistence-area relationships for the main analysis and different modifications of seed survival or seed germination rates. The curves shown are the average outcomes of 100 replicates for the random variation treatments, and the single curves obtained with the increase or decrease of each rate. Points and error bars have the same meaning as in Fig.2 and Fig. 4: average values across plot combinations, and standard errors of these averages.

### Supplementary Note 3: Local stability

Feasibility is, arguably, the most important condition for multispecies coexistence, as it evaluates whether there exists at least one equilibrium point in the system in which all species have positive abundances, i.e. no species goes extinct. These feasible equilibrium point(s) may, in turn, be locally stable in regards to perturbations in equilibrium abundances or not. We evaluated in our main analyses the feasibility of our sampled communities as a proxy for coexistence relationships, and here we analyze the local stability patterns, to understand whether feasibility and local stability are linked in our system.

We calculated local stability of the same species combinations, at the plot level, as in the feasibility analyses. For doing so, we calculated the steady-state solution of the equivalent Lotka-Volterra system with growth rates given by Supplementary equation 5 and the estimated interaction coefficients. We obtained the eigenvalues of the associated Jacobian matrix for each combination; the combination is locally stable if, in a discrete-time set of equations like this one, the maximum absolute value of the eigenvalues is $\leq 1$. That is, all eigenvalues are bounded below, or equal to, 1.

In our observed communities, almost no combination of species is locally stable (only 2 out of 471970 combining the homogeneous and heterogeneous parameterization, see associated code at `https://github.com/garciacallejas/SAR_coexistence`, in particular the scripts and results with prefix `S5`). Therefore, both properties are largely disconnected, and this analysis clearly suggests that the spatial arrangement of interactions in our system promotes feasibility over local stability, as observed in a similar analysis over the assembly of a seasonal plant-pollinator community (Saavedra et al. 2016, referenced in main text). Whether this is a general feature of seasonal systems or ecological communities in general remains to be explored. We argue that the lack of local stability is to be expected in annual systems that furthermore are modelled as closed communities, like in this study. Incorporating spatial dynamics in further studies will provide a more complete picture of the local stability of the system at the meta-community scale.
